# Ultrasonic local anesthetic uncaging enables noninvasive, targeted, and transient knock-out of brain function

**DOI:** 10.64898/2026.10.27.762226

**Authors:** Payton J. Martinez, Yun Xiang, Kanchan Sinha Roy, Martin Loynaz Prieto, Mahaveer P. Purohit, Brenda J. Yu, Alex R. Hart, Matine M. Azadian, Tenzing Kalsang, Lily Amaturo, Eujeong Choi, Rishabh Ranjan, Ruijia Sun, Kim Butts Pauly, Raag D. Airan

**Affiliations:** Department of Radiology, Stanford University, Stanford, CA, USA; Department of Biology, Stanford University, Stanford, CA, USA; Department of Materials Science & Engineering, Stanford University, Stanford, CA, USA; Department of Psychiatry & Behavioral Sciences, Stanford University, Stanford, CA, USA

## Abstract

Noninvasive, spatially localized brain inhibition would enable precise neural circuit interrogation, mapping, and treatment. We previously presented SonoRopi, an acoustically activatable liposomal formulation of ropivacaine, and demonstrated its utility for targeted peripheral nerve anesthesia. Here, we extend SonoRopi to localized brain anesthesia following intravenous administration and transcranial focused ultrasound application. Ultrasonic uncaging of SonoRopi releases ropivacaine, which crosses the intact blood-brain barrier to anesthetize the sonicated region. SonoRopi uncaging in mouse primary visual cortex suppressed visually evoked cortical activity for one hour, while uncaging in rat somatosensory hindlimb cortex produced lateralized, reversible hindlimb sensory anesthesia. Off-target control sonication preserved function in both paradigms. No acute histologic injury or open-field behavioral abnormalities were detected following this intervention. SonoRopi thereby enables noninvasive, spatially targeted, reversible brain inhibition with physiologic and behavioral efficacy in two rodent species.

## Introduction

Causal neural circuit analysis in either the healthy or diseased brain has relied upon the availability of natural permanent brain lesions post stroke or trauma, or induced temporary brain lesions via brain stimulation, opto-/chemogenetic modulation, or pharmacologic inactivation (*1–10*). The ideal tool for causal mapping of human brain function would be: noninvasive, spatiotemporally precise, able to target any brain region, have a reliable mechanism of action for the induced inactivation, be implemented without permanent change or injury to the underlying brain, could be paired with imaging for targeting and functional readout, and have low regulatory or physical barriers for clinical utilization.

Direct local delivery of anesthetics has been used preclinically and clinically to create reversible functional brain inactivation, or pharmacologic test lesions, in defined brain structures. Direct injection of agents like lidocaine, muscimol or tetrodotoxin into animal cortex has demonstrated the spatial spread and time course of cortical inactivation, establishing that local anesthetic blockade can suppress neural activity in spatially delimited brain regions over experimentally useful time windows, thereby becoming a mainstay of preclinical neuroscience (*2–5*). Clinically, intra-carotid infusion of anesthetics like amobarbital and propofol has long been used to determine lateralization of certain brain functions in the Wada test (*9*, *10*). These studies established the causal value of localized pharmacologic anesthetic inactivation, but the invasiveness of infusion via an injection cannula or needle has limited the clinical utilization of such studies.

A noninvasive pharmacologic test lesion technique would be valuable for experimental neuroscience, clinical target validation, and clinical treatment. Focused ultrasound (FUS) is well-suited for such a tool as it can noninvasively target acoustic energy to millimeter-scale brain regions and can be paired with drug carriers whose payload is released at the ultrasound focus. Prior work has established ultrasonic drug uncaging for targeted pharmacologic inactivation: perfluorocarbon nanoemulsions released propofol after FUS application to modulate cortical and subcortical activity with seconds-scale temporal control (*11*) and mapped whole-brain functional networks (*12*). Related platforms generalized ultrasound-activated particles for hydrophobic drug delivery, described focal pharmacologic neurointervention as a brain-targeting framework, showed that different anesthetic-loaded particles can produce targeted neuromodulation after transcranial FUS, and demonstrated ultrasound-triggered local anesthesia in peripheral tissues (*13–17*). Together, these studies demonstrate that uncaging of drug-loaded ultrasound-sensitive particles can define where a pharmacologic effect occurs. However, the drugs and vehicles used in those studies render translational challenges for use for clinical brain applications. Perfluorocarbon systems generally yield low drug loading, require complex manufacturing processes, and have stability profiles necessitating fresh manufacturing or re-activation immediately prior to administration. Gas-containing vehicles render risks of instability and damaging cavitation. Meanwhile, drugs like propofol render too short inactivation time periods for clinical utility, while drugs like muscimol or tetrodotoxin present regulatory barriers due to lack of prior clinical approvals. Additionally, some of the presented vehicles were designed around and validated with ultrasound parameters that render translational challenges for human transcranial application.

Recently, we developed acoustically activatable liposomes (AALs) with sucrose-containing aqueous cores that enhance ultrasound-triggered release without requiring gas cores or perfluorocarbon vaporization (*18*). This AAL platform enables site-targeted drug delivery and noninvasive neuromodulation and can encapsulate common clinically-utilized sodium-channel blocker local anesthetics including lidocaine, bupivacaine, and ropivacaine that partition intracellularly to produce longer inactivation times than GABAergic agents like propofol or muscimol (*18–21*). While we previously validated that local anesthetic uncaging with AALs can enable targeted peripheral nerve anesthesia (*18*), we note that local anesthetics like ropivacaine can also cross the intact blood-brain barrier to potentially enable targeted brain regional anesthesia suitable for causal neural circuit analysis. Additionally, AALs are efficaciously activated by the submegahertz ultrasound frequencies common to clinical transcranial FUS systems. These features make AALs well matched to the local brain inactivation problem: encapsulation can reduce nonspecific tissue exposure, while FUS can convert a circulating intravascular reservoir into focal release of a local anesthetic drug that can internalize intracellularly to inactivate neural tissue for minutes to hours. Additionally, AALs are constructed with validated pharmaceutical excipients and by common pharmaceutical manufacturing methods, lowering barriers to translation.

Here, we asked whether ropivacaine-loaded AALs, termed SonoRopi, could provide noninvasive, spatially targeted local anesthetic neuromodulation in the brain. We hypothesized that AAL encapsulation would reduce nonspecific ropivacaine exposure while allowing FUS to define where ropivacaine acts. We evaluated SonoRopi across formulation, pharmacokinetic, neurochemical, functional imaging, behavioral, and safety endpoints. We first characterized ultrasound-triggered ropivacaine release under flow conditions and assessed formulation stability. We then compared systemic biodistribution and blood pharmacokinetics of SonoRopi and free ropivacaine and used solid-phase microextraction (SPME) coupled to liquid chromatography-tandem mass spectrometry (LC-MS/MS) to measure ropivacaine pharmacodistribution after ultrasonic uncaging. To assess the ability of this system to induce a targeted functional test lesion, we targeted SonoRopi uncaging to the mouse primary visual cortex during widefield GCaMP imaging of visually evoked responses and separately targeted rat somatosensory hindlimb cortex during mechanical and thermal sensory testing. Finally, we assessed histological and open-field behavioral safety measures after brain-targeted ropivacaine uncaging. Together, these studies establish in two rodent species that SonoRopi uncaging enables a reversible pharmacologic brain test lesion suitable for causal neural circuit analysis.

## Results

### SonoRopi enables mechanical index-scaled ultrasonic ropivacaine uncaging

Following our prior work (*18*), SonoRopi had a mean hydrodynamic diameter of 166.71 +/- 3.8 nm, a polydispersity index (PDI) of 0.08 +/- 0.01, and a zeta potential of -49.2 mV, with ropivacaine loading of 4.51 +/- 0.42 mg/mL and 4.85 +/- 1.07% free drug. Cryo-electron microscopy (cryo-EM) showed that SonoRopi retained spherical vesicle morphology and visible bilayer structure after sonication, without gross aggregation, fusion, fractionation, or gaseous void formation (Fig. 1A-C). There was a downward shift in the particle size distribution after FUS exposure: the modal diameter shifted from 0.21 μm without FUS to 0.18 μm after FUS, and the histogram-weighted mean diameter decreased from 0.22 to 0.19 μm. These findings extend our prior observation that acoustically activatable liposomes decrease in size after sonication while retaining vesicular morphology, consistent with ultrasound-induced increase in drug membrane permeability and not a burst release. Notably the lack of visible gas elements with or without FUS indicates limited risk of damaging inertial cavitation with SonoRopi uncaging.

**Fig. 1.**
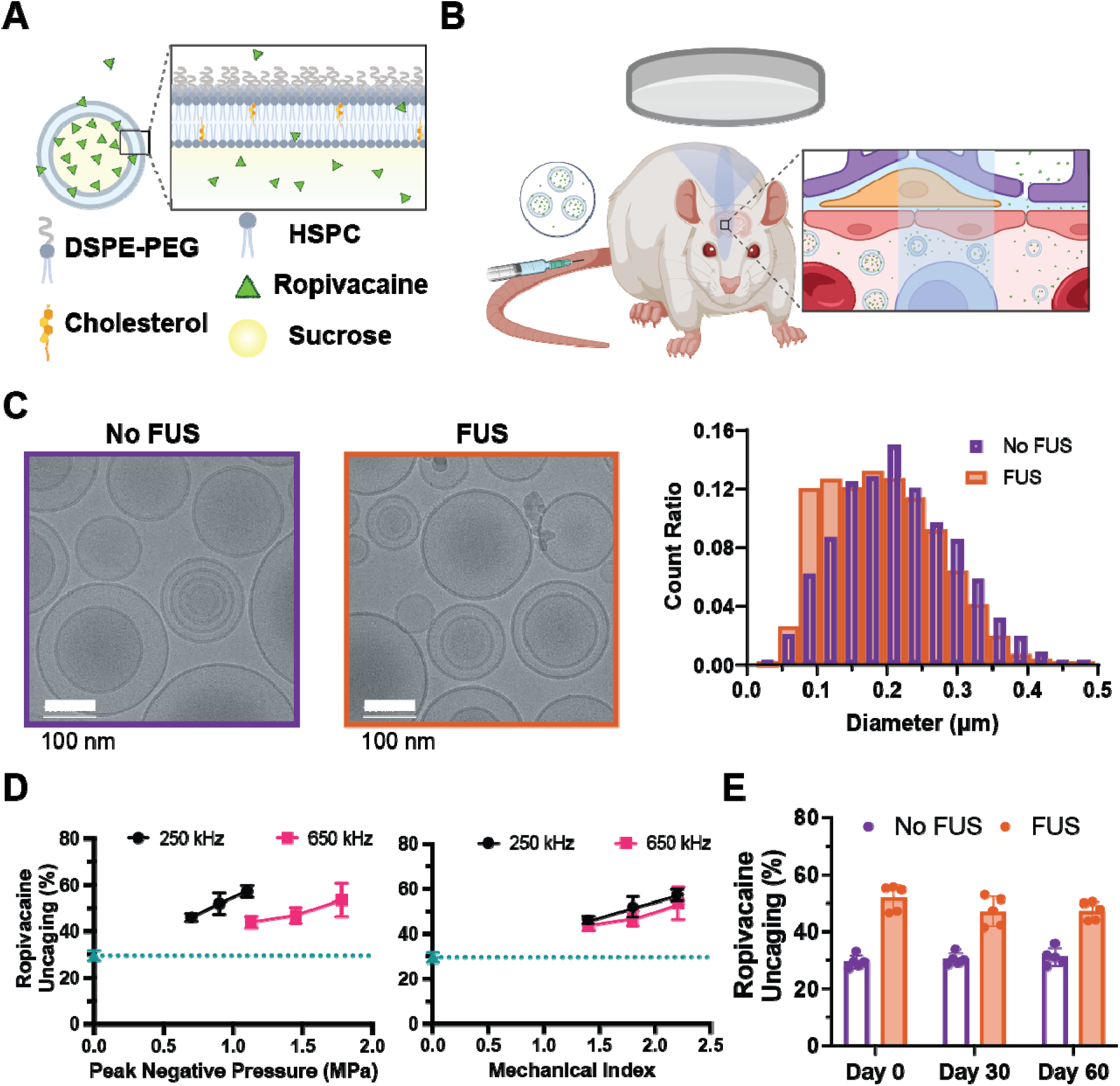
Characterization and ultrasound-triggered release of SonoRopi. (A) Schematic of SonoRopi, a ropivacaine-loaded acoustically activatable liposome composed of phospholipids, cholesterol, and DSPE-PEG, with 5% sucrose in the aqueous core. (B) Schematic of intravenous SonoRopi administration and focused ultrasound (FUS)-triggered ropivacaine release at th targeted brain site. (C) Representative cryo-electron microscopy (cryo-EM) images of SonoRopi with and without FUS exposure, with particle diameter distributions quantified from cryo-EM images and shown as count ratios. The modal particle diameter shifted from 0.21 μm without FUS to 0.18 μm after FUS, and the histogram-weighted mean diameter decreased from 0.22 to 0.19 μm. Scale bars, 100 nm. (D) Ultrasound-triggered ropivacaine release under plasma-flow conditions at 250 and 650 kHz, plotted as a function of peak negative pressure (left) and mechanical index (MI, right). Matched no-FUS flow samples were used to quantify the assay floor, indicated by the green dotted line. Data are mean ± SD; n = 4 independent release samples per condition. (E) Storage stability of SonoRopi at 4 °C, showing preserved FUS-triggered ropivacaine release through Day 60. Data are mean ± SD; n = 5 independent samples per storage time point; one-way ANOVA.

To quantify ultrasound-triggered ropivacaine release under conditions mimicking the intravascular environment, SonoRopi was diluted in plasma and passed through a FUS flow exposure system. FUS exposure produced a pressure-dependent increase in ropivacaine release at both 250 kHz and 650 kHz (Fig. 1D). At 250 kHz, release increased from 46.0 +/- 1.8% at 0.7 MPa to 57.3 +/- 2.6% at 1.1 MPa, exceeding the matched sham-flow floor measured under the same plasma-flow conditions (29.5 +/- 2.2%). A similar monotonic increase was observed at 650 kHz, with release rising from 43.9 +/- 2.6% at 1.13 MPa to 53.6 +/- 7.2% at 1.9 MPa. When this release profile was normalized by mechanical index (MI), the 250 kHz and 650 kHz datasets followed a comparable trend showing that SonoRopi acoustic responsiveness scaled with MI across the tested frequencies rather than rising with frequency, indicating a mechanical oscillation-type mechanism.

Towards clinical practicality, we evaluated whether SonoRopi retained acoustic responsiveness during extended refrigerated storage. Liposomes stored in refrigeration (2-5 °C) were assessed for ultrasound-triggered release at 1.1 MPa and 250 kHz after 2 months of storage (Fig. 1E). Release efficiency did not significantly differ across storage time points, indicating preserved ultrasound responsiveness over extended storage.

### Ultrasonic SonoRopi uncaging yields spatially selective ropivacaine delivery in vivo

To evaluate whether AAL encapsulation preserved circulating ropivacaine availability before local ultrasound activation, we first measured early blood pharmacokinetics after bolus administration of SonoRopi or dose-matched free ropivacaine, without FUS in rats (Fig. 2A). The pharmacokinetic curves separated immediately after dosing: SonoRopi reached a mean 1-min blood concentration of 151.69 ± 32.48 ng/100 μL, compared with 59.03 ± 13.04 ng/100 μL for free ropivacaine. Noncompartmental pharmacokinetic analysis supported a pronounced early-exposure difference between formulations (Table S1). SonoRopi had increased C_max_ compared with free ropivacaine (P = 0.0278), whereas AUC_0-t_ showed a strong trend that did not meet the nominal significance threshold (P = 0.0518). Other model-derived parameters, including MRT_0-t_, CL/D, and V_ss_/D, differed under the assumptions of the terminal log-linear analysis. This early circulating window informed the subsequent uncaging protocol, in which a 5-min SonoRopi infusion was paired with FUS delivered during the final 2.5 min to overlap sonication with circulating SonoRopi availability.

**Fig. 2.**
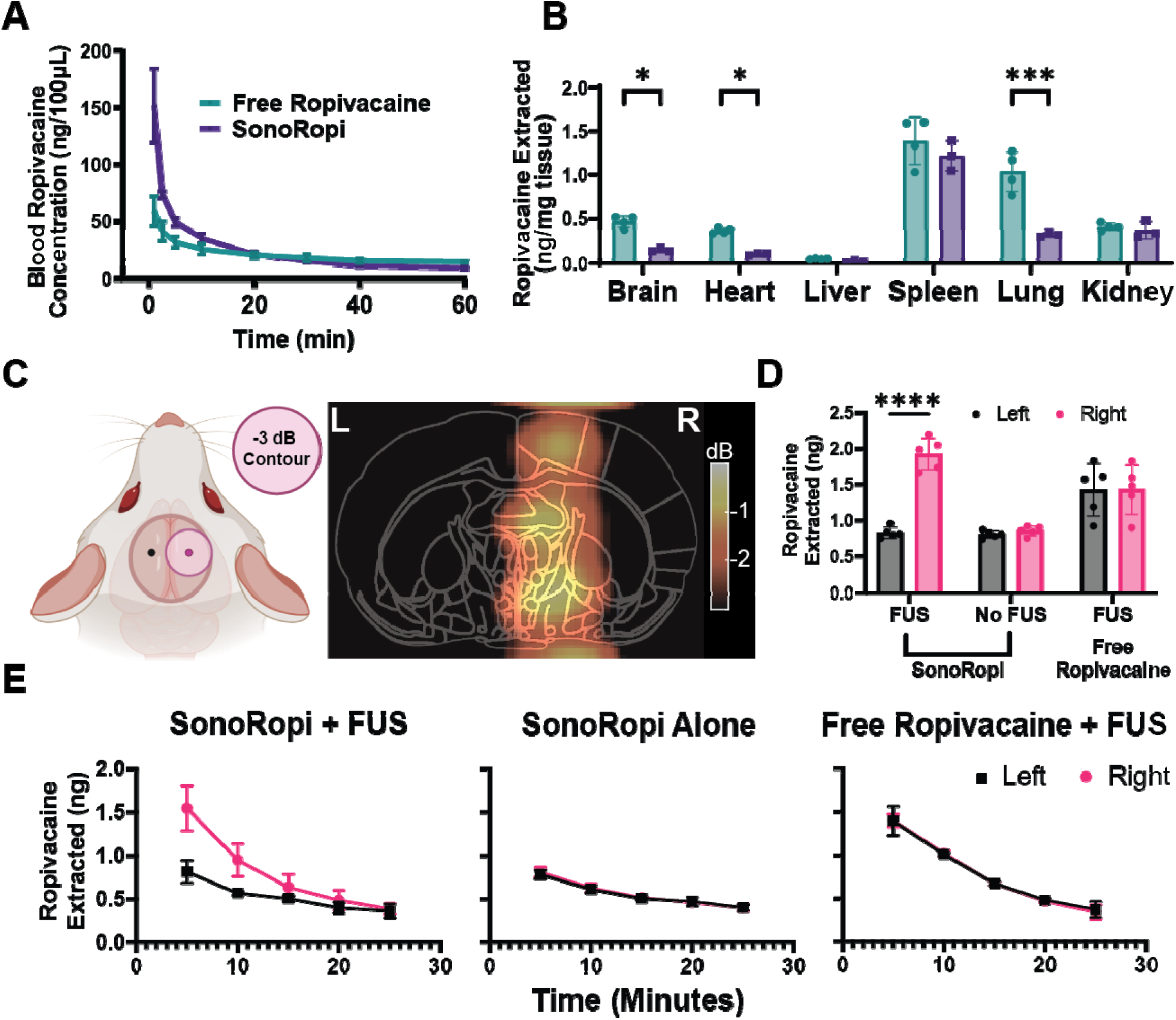
SonoRopi uncaging targets ropivacaine delivery while limiting off-target exposure. (A) Blood pharmacokinetics of ropivacaine after intravenous bolus administration of free ropivacaine or SonoRopi in the absence of FUS. n ≥ 3 rats per group. (B) Tissue ropivacaine concentrations after intravenous bolus administration of dose-matched free ropivacaine or SonoRopi analyzed by LC-MS/MS. n = 4 rats per group; two-way ANOVA with multiple comparisons between formulations within each tissue. (C) FUS targeting strategy and acoustic simulation (finite-element simulation of 250-kHz FUS beam with -3 dB focal contour) for right somatosensory hindlimb cortex (S1HL) delivery. (D) Hemisphere-specific ropivacaine extraction measured by solid-phase microextraction coupled to LC-MS/MS (SPME-LC-MS/MS) after SonoRopi + FUS, SonoRopi alone, or free ropivacaine + FUS. n = 5 rats per group; two-way repeated-measures ANOVA with treatment as the between-animal factor and hemisphere as the within-animal factor. (E) Time course of extracted ropivacaine after treatment. Left and right hemisphere SPME measurements show transient right-hemisphere enrichment after SonoRopi + FUS, whereas SonoRopi alone and free ropivacaine + FUS show identical left-right distribution and clearance. n = 4 rats per time point/group. Together, these measurements show that SonoRopi reduces off-target tissue exposure, maintains early circulating ropivacaine availability, and produces transient, FUS-dependent ropivacaine enrichment in the targeted hemisphere. Data are mean ± SD. *P < 0.05, ***P < 0.001 ****P < 0.0001. Exact P values are provided in the source data.

We then quantified tissue biodistribution following single-bolus intravenous administration of SonoRopi or dose-matched free ropivacaine using LC-MS/MS in rats (Fig. 2B). AAL encapsulation substantially reduced off-target accumulation across multiple organs. Free ropivacaine reached significantly higher concentrations than SonoRopi in the brain (0.47 ± 0.07 vs. 0.15 ± 0.02 ng/mg), heart (0.36 ± 0.03 vs. 0.10 ± 0.01 ng/mg), and lung (1.04 ± 0.22 vs. 0.32 ± 0.02 ng/mg), whereas spleen and kidney levels were more comparable between formulations, presumably reflecting their reticuloendothelial cell content. Notably, the lower brain and heart exposure after SonoRopi administration confirms reduced ropivacaine delivery to organs associated with systemic local-anesthetic toxicity (LAST) and therefore reduced risk of LAST with SonoRopi (*22*).

To verify the spatial precision of ultrasonic uncaging, we sampled the rat brain ropivacaine concentration with SPME-LC-MS/MS following SonoRopi or free ropivacaine administration, with and without FUS activation. First, finite-element simulations of the 250-kHz, 1.1-MPa sonication protocol confirmed the approach to align the -3 dB focal zone with the target right somatosensory hindlimb cortex (S1HL). These coordinates were used to guide bilateral placement of SPME probes via stereotaxic burr holes (Fig. 2C). SPME-LC-MS/MS revealed a pronounced, hemisphere-specific increase in ropivacaine extraction only when SonoRopi was combined with FUS (Fig. 2D). In the SonoRopi + FUS uncaging group extracted ropivacaine was higher in the sonicated right hemisphere than in the contralateral hemisphere (1.93 ± 0.22 ng vs. 0.83 ± 0.08 ng). In contrast, SonoRopi administered without FUS produced low, symmetric concentrations between hemispheres (0.86 ± 0.07 ng vs. 0.81 ± 0.05 ng), which likely is an overestimate of the true brain uptake as it reflects a floor effect due to concomitant blood sampling during the SPME probe insertion (see Fig. S1 of (*18*)). Meanwhile, free ropivacaine + FUS yielded higher bilateral brain levels that were symmetrically equal (1.43 ± 0.35 ng vs. 1.43 ± 0.37 ng). Importantly, contralateral ropivacaine exposure after SonoRopi + FUS remained below free-drug exposure, indicating that FUS increased drug levels at the target site without producing a global increase throughout the brain.

Time-course SPME measurements demonstrated that the SonoRopi uncaging-mediated local ropivacaine enrichment was transient (Fig. 2E). After SonoRopi uncaging, the right-left separation was largest at early sampling times and declined over the post-treatment window, converging toward control conditions by approximately 25 min. In contrast, SonoRopi alone showed low and symmetric left-right measurements over time, and free ropivacaine + FUS produced overlapping left and right hemisphere curves without a sonication-dependent separation.

### Ultrasonic SonoRopi uncaging suppresses visually evoked cortical activity in mice

To determine whether this targeted pharmacodistribution enabled by SonoRopi uncaging (Fig. 2) would result in the localized suppression of cortical activity necessary for a test lesion, we performed widefield calcium imaging of visually evoked responses in Thy1-GCaMP6s mice, which express the calcium-sensitive fluorescent protein GCaMP6s in excitatory neurons throughout the cortex (*23*). After recording baseline visually evoked activity, mice received one of four treatments: FUS targeted to the right primary visual cortex (V1) alone, SonoRopi alone, SonoRopi + FUS targeted to right V1, or SonoRopi + FUS targeted to an off-target left medial prefrontal cortex (mPFC) site (Fig. 3A,B). SonoRopi was administered intravenously at a ropivacaine-equivalent dose of 2.5 mg/kg, and FUS was delivered at 250 kHz, 1.1 MPa estimated in situ peak negative pressure, and 25% duty cycle. Visual responses were evoked by flash stimulation of the contralateral eye, and imaging sessions were repeated at 5 (immediate), 30, and 60 min after treatment.

**Fig. 3.**
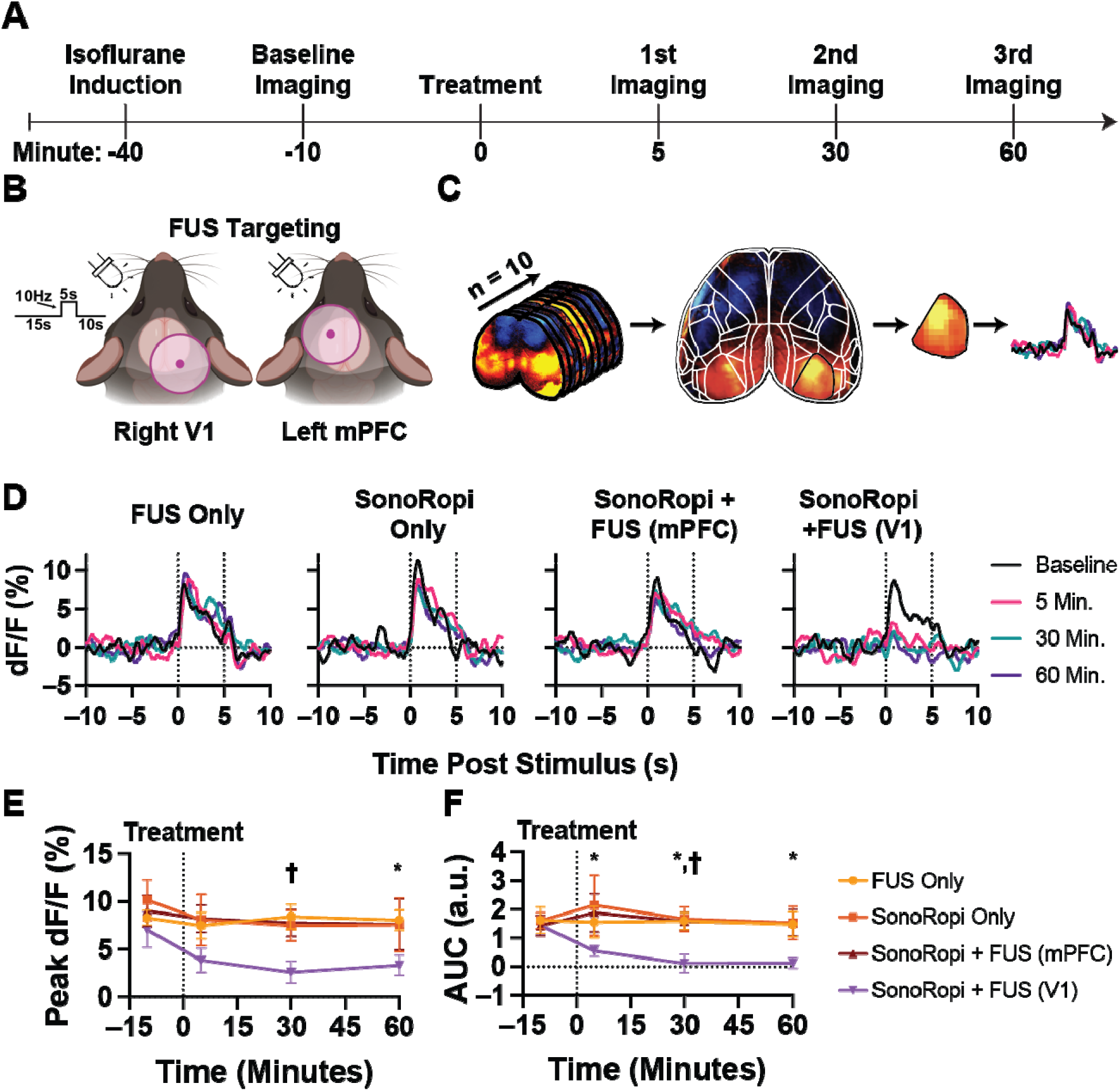
SonoRopi uncaging in V1 suppresses visually evoked cortical activity. (A) Experimental timeline. Adult Thy1-GCaMP6s mice were induced with isoflurane at -40 min, imaged at baseline at -10 min, treated at 0 min, and re-imaged at 5, 30, and 60 min after treatment. (B) Visual stimulation and FUS targeting strategy. Each trial consisted of a 15-s pre-stimulus baseline, 5-s flash stimulation at 10 Hz, and 10-s post-stimulus recording, with FUS targeting either right primary visual cortex (V1) or left medial prefrontal cortex (mPFC). (C) Widefield dual-wavelength GCaMP imaging and processing workflow. Ten repeated visual stimulation trials per session. Calcium-dependent ΔF/F signals were extracted from manually defined right V1 regions of interest and aligned to the visual stimulation period. (D) Stimulus-locked ΔF/F traces from right V1 across treatment groups and imaging time points, showing visually evoked responses before and after treatment. (E) Peak and (F) area under the curv (AUC) visually evoked ΔF/F during the 5-s visual stimulation period. For E and F, data are mean ± SEM; n = 6 mice per group. Baseline responses were compared across groups using one-way ANOVA (P = 0.97). Treatment effects across time and group were analyzed using repeated-measures two-way ANOVA with Tukey correction for multiple comparisons. *P < 0.05 for SonoRopi + FUS targeted to V1 compared with off-target SonoRopi + FUS targeted to mPFC at the indicated time point; †P < 0.05 for SonoRopi + FUS targeted to V1 compared with its own baseline. Together, these measurements show that on-target SonoRopi + FUS transiently suppresses visually evoked V1 activity, whereas off-target mPFC sonication does not produce comparable V1 suppression.

Widefield calcium signals were processed using a dual-wavelength correction pipeline, in which 470 nm excitation was used to measure calcium-dependent fluorescence from the genetically encoded calcium indicator GCaMP and 405 nm excitation was used as a reference channel to correct for hemodynamic and motion-related artifacts (Fig. 3C). Stimulus-locked changes in fluorescence (ΔF/F) were calculated relative to the mean pre-stimulus fluorescence for each trial and extracted from manually defined right V1 regions of interest. Pretreatment visually evoked responses were comparable across treatment groups, indicating similar cortical activation before intervention. Because isoflurane can attenuate visually evoked cortical responses at higher concentrations, we first evaluated the effect of maintenance isoflurane on flash-evoked widefield calcium responses. Maintenance at 1% isoflurane produced minimal deviation from the baseline response, whereas higher isoflurane concentrations produced larger response deviations (Fig. S1). We therefore used 1% isoflurane for maintenance sedation for all visually evoked response imaging experiments.

Representative cortical activity maps showing visually evoked responses at baseline and 30 min post-treatment further are shown in Fig. S2. Stimulus-locked right V1 traces, normalized to the mean pre-stimulus fluorescence for each trial, showed robust visually evoked responses at the pretreatment baseline across groups. After treatment, the evoked response was markedly reduced only in mice receiving SonoRopi + FUS targeted to right V1 (Fig. 3D). In contrast, mice treated with right V1 FUS alone, SonoRopi alone, or SonoRopi + FUS targeted to the off-target left mPFC site retained visually evoked V1 responses with no difference from the baseline condition. Quantitation confirmed a reduction in visually evoked cortical activity in the SonoRopi + FUS V1 group only (Fig. 3E,F). Peak ΔF/F decreased from 7 ± 4% at baseline to 4 ± 3% at 5 min and reached a nadir of 2 ± 2% at 30 min after treatment, with minimal recovery to 3 ± 2% at 60 min. The area under the stimulus-evoked ΔF/F curve showed a similar temporal profile, decreasing from 1.44 ± 0.76% at baseline to 0.55 ± 0.44% at 5 min, reaching 0.08 ± 0.45% at 30 min, and minimal recovery to 0.13 ± 0.42% at 60 min. At the indicated time points, on-target SonoRopi + FUS in V1 differed from off-target SonoRopi + FUS in mPFC and from its own baseline, supporting focal rather than systemic suppression of visually evoked activity. Together, these data indicate that V1 cortical activity suppression with SonoRopi uncaging required both SonoRopi and on-target spatially-selective FUS delivery, and was not produced by ultrasound exposure alone (potentially reflecting that this ultrasound protocol is optimized for uncaging rather than neuromodulation (*24*)), circulating SonoRopi alone, or FUS-triggered release at a distant cortical site and subsequent transport of the freed ropivacaine to the visual cortex.

### SonoRopi uncaging produces lateralized and reversible sensory modulation in rats

To determine whether focal ropivacaine uncaging could yield a behavioral modulation akin to the observed physiologic modulation (Fig. 3), and to assess its efficacy in another species, we targeted FUS to the right somatosensory hindlimb cortex (S1HL) in adult rats receiving FUS alone, SonoRopi alone, or combined SonoRopi + FUS (Fig. 4A, D). Because right S1HL represents the contralateral left hindlimb, sensory responses were analyzed separately in the left and right hindlimbs. Mechanical sensitivity was assessed using von Frey testing, and thermal sensitivity was assessed using a 50 °C hot plate assay at baseline and after treatment.

**Fig. 4.**
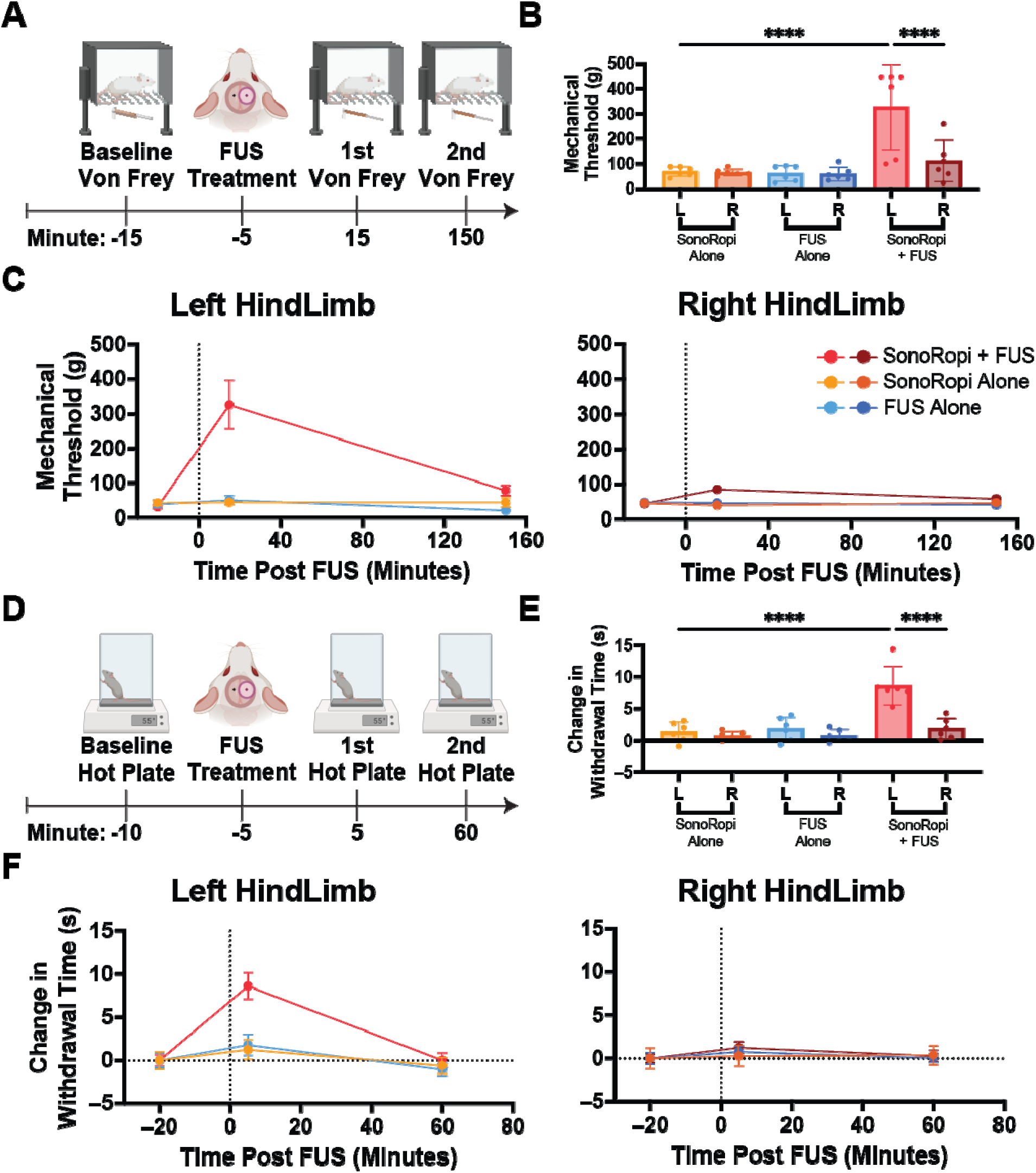
SonoRopi uncaging in S1HL produces lateralized and reversible sensory anesthesia. (A) Experimental timeline. Baseline von Frey testing was performed in Sprague Dawley rats at - 15 min, FUS treatment was delivered at -5 min, and post-treatment von Frey testing was performed at 15 and 150 min. (B) Peak mechanical withdrawal thresholds for left and right hindlimbs across treatment groups after right S1HL targeting. (C) Time course of mechanical withdrawal thresholds after treatment, shown separately for left and right hindlimbs. (D) Experimental timeline. Baseline 50 °C hot plate testing was performed at -10 min, FUS treatment was delivered at -5 min, and post-treatment hot plate testing was performed at 5 and 60 min. (E) Peak baseline-normalized changes in thermal withdrawal latency for left and right hindlimbs across treatment groups after right S1HL targeting. (F) Time course of baseline-normalized changes in thermal withdrawal latency after treatment, shown separately for left and right hindlimbs. For B to F, data are mean ± SD with individual animals overlaid where shown; n = 6 rats per group. Peak outcomes in B and E were analyzed using two-way repeated-measures ANOVA with treatment group as the between-animal factor and hindlimb side as the within-animal factor. Time-course analyses in C and F accounted for repeated measurements within animals. ****P < 0.0001 for indicated comparisons. Together, these measurements show that SonoRopi + FUS targeted to right S1HL increased mechanical thresholds and thermal latencies selectively in the contralateral left hindlimb, with recovery over time, supporting lateralized and reversible sensory modulation.

S1HL-targeted SonoRopi + right S1HL FUS produced a rapid and lateralized increase in mechanical withdrawal threshold. In the contralateral left hindlimb, mechanical withdrawal thresholds increased within 15 min after treatment, rising from a baseline of 47.8 ± 10.4 g to 326.8 ± 171 g (Fig. 4B,C). In contrast, left hindlimb thresholds remained near baseline in rats treated with SonoRopi alone (44.7 ± 18 g) or FUS alone (60 ± 31 g). Peak withdrawal thresholds were significantly higher in the left hindlimb of the SonoRopi + right S1HL FUS group than in all other treatment and limb conditions (*P* < 0.0001; Fig. 4B). The ipsilateral right hindlimb did not show a comparable increase, indicating that the sensory effect was lateralized to the limb represented by the targeted cortical region.

Thermal sensitivity showed a similar contralateral and reversible modulation. In the SonoRopi + right S1HL FUS group, hot plate withdrawal latency in the left hindlimb increased rapidly after treatment, reaching 8.6 ± 1.5 s above baseline at 5 min (Fig. 4E,F). By comparison, SonoRopi alone and FUS alone produced no significant changes in withdrawal latency (1.3 ± 1.1 s and 1.8 ± 1.2 s, respectively). Peak thermal responses were significantly greater in the contralateral left hindlimb of the SonoRopi + right S1HL FUS group than in all other treatment and limb conditions (*P* < 0.0001; Fig. 4E), whereas the ipsilateral right hindlimb remained unchanged.

Both mechanical and thermal effects were transient. Mechanical thresholds gradually returned toward baseline by the final von Frey time point at 150 min, and thermal withdrawal latencies returned toward baseline by the final hot plate time point at 60 min (Fig. 4C,F). Both mechanical and thermal behavioral effects were appropriately lateralized and only observed with both SonoRopi administration and on-target FUS, similar the physiologic effect in mice (Fig. 3).

### SonoRopi uncaging produces no histologic injury or open-field behavioral abnormalities

We next assessed whether the same brain-targeted uncaging paradigm produced detectable tissue injury or unexpected behavioral abnormalities. This safety experiment used the same right S1HL target and sonication parameters as the mechanical and thermal sensory testing in Fig. 4, allowing us to determine whether the sensory effects observed in Fig. 4 reflected reversible neuromodulation rather than irreversible injury to the targeted brain region or anesthesia of the adjacent motor cortex. Adult Sprague-Dawley rats received sham saline, FUS alone, or SonoRopi + right S1HL FUS, and brains were collected 72 h after treatment for histological analysis (Fig. 5A). Coronal sections spanning the ultrasound-targeted region were evaluated for tissue structural changes, neurodegeneration, microglial activation, and astrocytic activation.

**Fig. 5.**
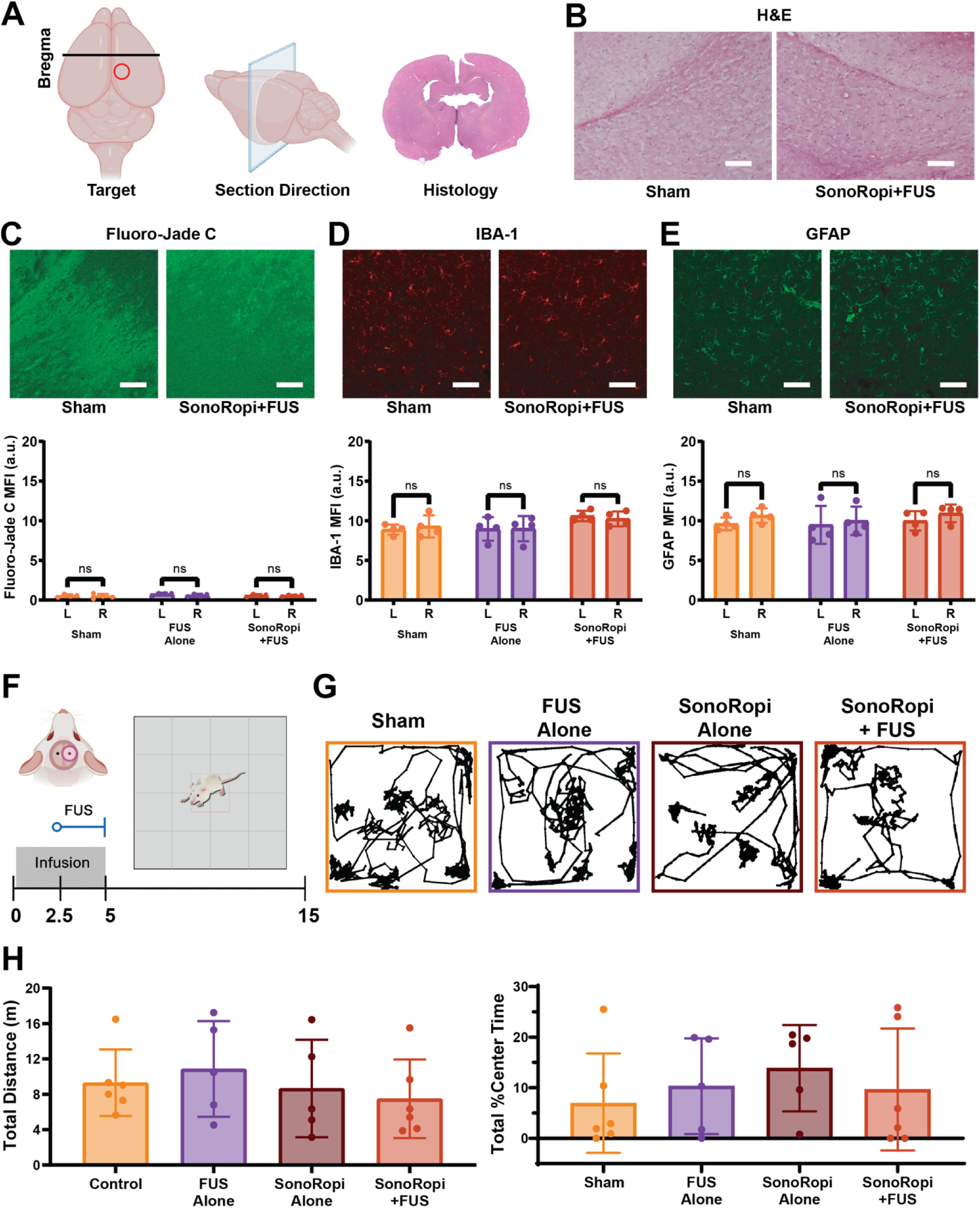
SonoRopi uncaging produces no histologic injury or open-field behavioral changes. (A) Schematic of histological safety assessment in Sprague Dawley rats after right S1HL-targeted treatment using the same target and sonication parameters as the mechanical and thermal sensory testing in Fig. 4 indicating the ultrasound beam, sampled brain sections relative to bregma, and the histology region of interest. Representative hematoxylin and eosin (H&E; B), Fluoro-Jade C (C), IBA-1 (D), and GFAP (E) stained sections from sham and SonoRopi + FUS animals, showing preserved tissue morphology, and no degenerating neurons, activated microglia or astrocytes in the targeted region. Scale bar, 100 μm. (F) Experimental timeline for open-field behavioral assessment after right S1HL-targeted treatment. Rats received intravenous saline or SonoRopi with or without FUS during infusion and were recorded in the open-field arena after treatment. (G) Overlay open-field movement tracks for sham, FUS alone, SonoRopi alone, and SonoRopi + FUS groups. (H) Quantitative analysis of the tracks in G, showing open-field total distance traveled and percentage of time spent in the center across treatment groups. For C to E, absolute MFI data are mean ± SD; n = 4 rats per group; repeated-measures two-way ANOVA accounting for hemisphere within animal; ns, not significant for the indicated left-right comparisons. For H, data are mean ± SD with individual animals overlaid; n = 5-6 rats per group; one-way ANOVA for each open-field endpoint. Together, these histological and behavioral safety measurements show that right S1HL-targeted SonoRopi + FUS did not produce detectable histologic injury or open-field behavioral abnormalities under the tested conditions.

Hematoxylin and eosin (H&E) staining showed preserved tissue architecture in the targeted region across treatment groups, without gross parenchymal disruption, hemorrhage, edema, or overt cellular morphology changes (Fig. 5B). Fluoro-Jade C staining, which labels degenerating neurons, remained low across groups and did not show an obvious neurodegenerative signal after SonoRopi + right S1HL FUS compared with sham or FUS-alone controls (Fig. 5C). Ionized calcium-binding adaptor molecule 1 (IBA-1) immunostaining labeled resident microglia, and glial fibrillary acidic protein (GFAP) immunostaining labeled astrocytes. As expected, IBA-1 and GFAP signals were detectable in normal brain parenchyma, but neither marker showed an increase after FUS alone or SonoRopi + FUS relative to sham tissue. Quantification by absolute mean fluorescence intensity (MFI) showed no increase in IBA-1+ or GFAP+ signal after SonoRopi + FUS, indicating no detectable microglial activation or astrocytic activation or gliosis at the treated site under the tested conditions (Fig. 5D,E).

We also evaluated whether treatment produced acute behavioral abnormalities in an open-field assay. Rats received intravenous saline or SonoRopi infusion with or without right S1HL-targeted FUS were placed in an open-field arena 15 min after treatment and recorded from above for 15 min (Fig. 5F). Overlay movement tracks showed similar exploratory coverage across treatment groups, including sham, FUS alone, SonoRopi alone, and SonoRopi + FUS animals (Fig. 5G). Quantitative analysis of these tracks showed that total distance traveled remained in a similar range across groups, approximately 8 to 11 m during the recording period, and that the percentage of time spent in the center remained broadly comparable, approximately 7 to 14% across groups (Fig. 5H). Thus, SonoRopi + FUS did not produce detectable reductions in locomotor activity or center-time behavior under the tested conditions.

## Discussion

This study supports SonoRopi as an ultrasound-gated local anesthetic platform for noninvasive and spatially targeted neural inhibition suitable for inducing reversible test lesions for causal neural circuit analysis. SonoRopi encapsulation limits nonspecific ropivacaine tissue exposure and provides a circulating drug reservoir, so that FUS can define the site at which the active anesthetic is released. SonoRopi uncaging leads to focal brain ropivacaine enrichment, inducing transient cortical suppression and lateralized physiologic and behavioral modulation, without detectable injury or adverse functional effect. Towards clinical translatability, SonoRopi is produced solely with validated pharmaceutical excipients and with common pharmaceutical production methods and is verified here to have efficacy when activated by the sub-megahertz frequencies of clinical transcranial FUS systems.

For functional clinical neurologic and psychiatric applications, this study demonstrates that FUS can serve as a spatial gate for pharmacology, in addition to its uses as a direct neuromodulatory stimulus or for thermal ablation (*25*). Clinical FUS workflows, with similar ultrasound frequencies as used here, have established the feasibility of planning and delivering transcranial ultrasound with image-guidance, correction for skull absorption and aberration, and intra-procedural monitoring (*26*, *27*). AALs like SonoRopi add a drug-release dimension. The presented rodent experiments are proofs-of-concept of analogous clinical image-guided procedures that could integrate FUS with neuronavigation, individualized acoustic modeling, safety monitoring, and functional neuroimaging and electrophysiology (*26*, *28*) to enable targeted, reversible, and safe functional knock out of millimetric brain regions for functional brain mapping or adjunctive therapy.

We recognize that these proof-of-concept studies in healthy animals do not yet establish therapeutic efficacy or relevance in disease models. The SPME technique used for the in vivo brain pharmacodistribution studies (Fig. 2) provides evidence of local drug enrichment at the sonicated target, but does not define absolute extracellular ropivacaine concentration, cell-type-specific exposure, or the exact parenchymal distribution of released drug. As the SPME probes traverse blood vessels when inserted, the SPME data are also confounded by the blood concentration of ropivacaine in circulating SonoRopi, yielding a nonspecific floor effect in these data (see Fig. S1 of ref. (*18*)). Future translational studies would benefit from noninvasive or minimally invasive pharmacodynamic readouts, compared to the invasiveness of SPME. These functional studies were performed following uncaging at selected cortical targets; the spatial resolution, effective dose, and duration of inhibition may differ across subcortical vs cortical or white matter targets. Additionally, while histology and open-field testing showed no detectable injury or behavioral abnormalities at the tested time points, this single safety window does not define a complete safety and toxicologic analysis.

Towards translation, particular clinical protocols would benefit from established dose-response relationships across SonoRopi dose, ultrasound pressure, duty cycle, pulse duration, and total sonication time. For potential chronic applications, multi-dose treatment studies will be needed to determine whether repeated local anesthetic uncaging yields cumulative tissue injury, inflammatory activation, blood-brain barrier (BBB) disruption, or systemic toxicity. Finally, assessment in disease-relevant models may be needed to support application in hyperexcitable or excitation/inhibition imbalanced circuits like in focal epilepsy.

## Materials and Methods

### Experimental design

This study was designed to determine whether intravenously administered ropivacaine-loaded acoustically activatable liposomes (SonoRopi) could be triggered by focused ultrasound (FUS) to produce localized, reversible neuromodulation in the central nervous system while limiting systemic ropivacaine exposure. Experiments were organized around formulation characterization, in vitro ultrasound-triggered release, in vivo biodistribution and local brain uncaging, functional widefield calcium imaging, behavioral sensory testing, and histological safety assessment. The formulation and analytical workflows were adapted from our previously described acoustically activatable liposome platform (*18*).

For in vivo uncaging experiments, SonoRopi or control treatments were administered intravenously by a 5 min infusion, and FUS was applied during the final 2.5 min of the infusion unless otherwise specified. In rat brain experiments, FUS was targeted to the right somatosensory hindlimb cortex (S1HL), and sensory outcomes were evaluated primarily in the contralateral left hindlimb. In mouse visual cortex experiments, FUS was targeted to right primary visual cortex (V1) or to an off-target left medial prefrontal cortex (mPFC) control site. Animals were assigned to treatment groups according to the experimental timelines shown in the corresponding figures. Sample sizes, exclusion criteria, and randomization/blinding procedures are reported in the figure legends or in the Statistical analysis section where applicable.

### Animals

All animal procedures were approved by the Stanford University Administrative Panel on Laboratory Animal Care (APLAC protocol 32874). Male Sprague-Dawley rats (Charles River Laboratories), 12 to 24 weeks old and weighing 300 to 400 g, were maintained on a standard light-dark cycle. Adult Thy1-GCaMP6s GP4.3 transgenic mice (Jackson Labs stock #024275) expressing GCaMP6s in cortical excitatory neurons were used for widefield calcium imaging and maintained under a standard light-dark cycle.

### Materials

All reagents were used as received unless otherwise specified. Hydrogenated soy phosphatidylcholine (HSPC; Lipoid S PC-3) and DSPE-PEG2000 (PE 18:0/18:0 PEG 2000) were obtained from Lipoid. Cholesterol (PhytoChol Inject HU) was obtained from Evonik. Ropivacaine hydrochloride was obtained from AstaTech. Ammonium sulfate was obtained from HiMedia. Absolute ethanol, sucrose, sodium chloride, phosphate-buffered saline (PBS), HPLC-grade water, LC/MS-grade water, methanol, acetonitrile, 2-propanol, trifluoroacetic acid, and formic acid were obtained from Fisher Scientific unless otherwise specified. HEPES buffer solution and L-histidine monohydrochloride monohydrate were obtained from Sigma-Aldrich. Milli-Q water was used to prepare aqueous buffers.

Tangential flow filtration (TFF) filters were obtained from Repligen. Polycarbonate extrusion membranes and 220 nm PVDF sterile filters were obtained from Millipore. PD-10 Sephadex G-25 columns and Sephacryl S-500 resin were obtained from Cytiva. Ropivacaine USP reference standard was obtained from Sigma, and ropivacaine-d7 was obtained from Cayman Chemical. C18-coated BioSPME 96-pin devices were obtained from Millipore Sigma. Isoflurane was obtained from Dechra. Paraformaldehyde, hematoxylin and eosin staining reagents, Fluoro-Jade C reagents, anti-GFAP antibody, anti-IBA-1 antibody, fluorescent secondary antibodies, and nuclear counterstain were obtained from Antibodies Inc. and Abcam. Von Frey filaments, elevated mesh testing chambers, and the 50°C hot-plate apparatus were obtained from Bioseb.

### Focused ultrasound system and calibration

The custom FUS transducer used for in vivo studies was constructed using a 100-mm-diameter, 100-mm-focal-distance piezoelectric element obtained from Piezo Hannas (Wuhan) Tech Co., Ltd. The transducer had a center frequency of 250 kHz. Acoustic coupling to the animal’s head was achieved using a water-filled coupling cone sealed with a Mylar membrane, with ultrasound gel applied at the membrane–scalp interface. Waveforms were generated using a Siglent SDG2042X function generator and amplified with a 2100L RF power amplifier (Electronics & Innovation, Ltd., Rochester, NY, USA) before being delivered to the transducer. Acoustic output was measured using an annually calibrated Onda HGL-series hydrophone and recorded with a Keysight DSOX1204G oscilloscope.

### Preparation of SonoRopi

Ropivacaine-loaded acoustically activatable liposomes, hereafter referred to as SonoRopi, were prepared by adapting our previously described sucrose-containing acoustically activatable liposome platform (*18*). Briefly, HSPC, cholesterol, and DSPE-PEG2000 were dissolved in heated ethanol at a molar ratio of 52.8:42.3:4.8. The lipid solution was diluted to 10% ethanol with an aqueous internal buffer containing 250 mM ammonium sulfate and 5% sucrose to form multilamellar vesicles. The suspension was then extruded ten times through 200 nm polycarbonate membranes at 65 to 70 degrees C to generate nanoscale liposomes.

An initial TFF step was performed against 10 mM HEPES and 145 mM NaCl, pH 7.4, to remove ethanol and external ammonium sulfate while establishing a transmembrane ammonium sulfate gradient for active drug loading. Ropivacaine hydrochloride was then added to the liposome suspension at a final concentration of 10 mg/mL and incubated at 55 degrees C for 1.5 h to enable remote loading into the liposomal core. After loading, a second TFF step was performed against 10 mM histidine buffer, pH 7.4, containing sucrose for osmotic balance. This second TFF step served to remove unencapsulated ropivacaine, to switch the external buffer to the long term storage buffer and to concentrate the final liposome suspension to the desired drug and lipid concentrations. The final SonoRopi suspension was sterile-filtered through a 220 nm PVDF filter and stored at 4 degrees C until use. The 5% sucrose internal formulation was selected based on prior work showing that sucrose incorporation increases ultrasound-triggered drug uncaging while preserving formulation stability, including in ropivacaine-loaded liposomes (*18*).

### Particle size and morphology characterization

SonoRopi particle size distribution, polydispersity index (PDI), and zeta potential were measured by dynamic light scattering (DLS) using a Malvern Zetasizer Nano ZS90, as previously described for acoustically activatable liposomes (*18*). Measurements were performed on purified liposome suspensions before FUS exposure and, where indicated, after storage at 4°C. Liposome morphology was evaluated by cryo-electron microscopy (cryo-EM) using purified SonoRopi samples before and after sonication. Cryo-EM images were assessed for vesicle shape, bilayer morphology, aggregation, fusion, and void-like or bubble-like gas structures, and particle diameters were quantified from cryo-EM images as described in the Supplementary Materials.

### HPLC quantification of ropivacaine

Ropivacaine concentrations in formulation and in vitro release samples were quantified by high-performance liquid chromatography (HPLC) using an Agilent 1260 Infinity II HPLC system with UV detection, as adapted from our prior AAL analytical workflow (*18*). Chromatographic conditions, gradient elution, and unencapsulated drug quantification procedures are provided in the Supplementary Materials (Table S2).

For total ropivacaine quantification, liposome samples were disrupted with methanol before HPLC analysis. For unencapsulated ropivacaine quantification, liposome suspensions were separated from free ropivacaine using PD-10 size-exclusion chromatography, and ropivacaine in the liposome-containing and free-drug fractions was quantified by HPLC as described in the Supplementary Materials.

### In vitro ultrasound-triggered release assay

Ultrasound-triggered release of ropivacaine from SonoRopi was measured using a flow-through FUS exposure design adapted from our previously described acoustically activatable liposome release assay (*18*). Briefly, SonoRopi was diluted 30-fold in canine plasma to mimic the protein-containing intravascular environment and passed through tubing positioned across the focal zone of either a 250 kHz or 650 kHz FUS transducer at a flow rate of 130 μL/min. Samples were exposed to FUS at the indicated peak negative pressures and 25% duty cycle while flowing through the acoustic focus; matched sham-flow samples were passed through the same flow path without ultrasound exposure to quantify passive leakage.

After sonication, released ropivacaine was separated from liposome-associated ropivacaine by size-exclusion chromatography using Sephacryl S-500 resin, which separates nanoscale liposomes and proteins in the plasma from free drug based on the size cutoff, as previously described for acoustically activatable liposomes (*18*). The liposome-containing fraction and free-drug fraction were collected separately, and ropivacaine in each fraction was quantified by HPLC. Ultrasound-triggered release was calculated as the amount of ropivacaine in the free-drug fraction divided by the sum of ropivacaine in the free-drug and liposome-containing fractions. Release was assessed across pressures at both 250 kHz and 650 kHz and plotted as a function of peak negative pressure and mechanical index. For storage stability studies, SonoRopi stored at 4 degrees C was evaluated using the same release assay at the indicated time points.

### LC-MS/MS analysis of ropivacaine in biological samples

Ropivacaine concentrations in blood, tissue homogenates, and SPME extracts were quantified by liquid chromatography-tandem mass spectrometry (LC-MS/MS) using an Agilent 1290 Infinity LC system coupled to an Agilent 6490 triple quadrupole LC/MS system with iFunnel technology and Agilent Jet Stream electrospray ionization operated in positive-ion mode. Biological samples were prepared by protein precipitation with cold acetonitrile containing ropivacaine-d7 internal standard, and SPME probes were desorbed in methanol/water (9:1, v/v) containing 1% formic acid and internal standard. Detailed source parameters, MRM transitions, chromatographic conditions, and calibration procedures are provided in the Supplementary Materials (Tables S3 to S5).

LC-MS/MS chromatographic separation was performed on an Agilent ZORBAX RRHD Eclipse Plus C18 column with a matching guard column using water and acetonitrile mobile phases containing 0.3% formic acid. Ropivacaine was quantified using multiple-reaction monitoring transitions for ropivacaine and ropivacaine-d7. Tissue concentrations were normalized to tissue mass and reported as ng/mg tissue, blood concentrations were reported as ng/100 μL blood unless otherwise indicated, and SPME measurements were reported as ng extracted per probe.

### Rat biodistribution and pharmacokinetic analysis

Ropivacaine biodistribution and pharmacokinetics were assessed in adult Sprague-Dawley rats using a dose-matched design adapted from our previously described in vivo acoustically activatable liposome studies (*18*). For biodistribution and blood pharmacokinetic analyses, free ropivacaine and SonoRopi were administered intravenously by bolus injection at ropivacaine-equivalent doses of 2 mg/kg. For biodistribution analysis, rats were euthanized 1 h after administration and transcardially perfused with 1x PBS to remove circulating blood. Brain, heart, liver, lung, spleen, kidney, spinal cord, and other indicated tissues were collected, weighed, homogenized, and processed for LC-MS/MS analysis. Ropivacaine concentrations were normalized to tissue mass and reported as ng ropivacaine per mg tissue. For pharmacokinetic analysis, venous blood samples were collected into EDTA at the indicated time points after free ropivacaine or SonoRopi administration, processed for LC-MS/MS analysis, and reported as ropivacaine concentration in blood or plasma.

### SPME measurement of FUS-triggered ropivacaine release in rat brain

Local FUS-triggered ropivacaine delivery in rat brain was quantified by solid-phase microextraction (SPME) coupled to LC-MS/MS using a procedure adapted from our prior AAL brain pharmacodistribution studies (*18*). Adult Sprague-Dawley rats received intravenous free ropivacaine, SonoRopi, or vehicle control at ropivacaine-equivalent doses of 2 mg/kg. Treatments were delivered by a 5 min intravenous infusion, with FUS or sham treatment applied unilaterally to the right S1HL during the final 2.5 min of infusion using a 250 kHz FUS transducer at an estimated in situ peak negative pressure of 1.1 MPa and a 25% duty cycle delivered as 50 ms pulses repeated every 200 ms. Hydrophone-informed acoustic modeling was used to align the -3 dB focal zone with the right S1HL and to define the contralateral non-sonicated control site.

After treatment, bilateral stereotaxic burr holes were made over the sonicated right S1HL and the contralateral control region. C18-coated BioSPME probes were mounted in a custom stereotaxic holder and inserted 3 mm ventral to the brain surface, positioning the absorptive coating approximately 2.5 mm below the cortical surface. Probes were left in contact with brain tissue for 5 min, removed, and briefly washed to minimize residual blood contamination. Extracted ropivacaine was desorbed from each probe in 50 microliter of MeOH/H2O (9:1, v/v) containing 1% formic acid and quantified by LC-MS/MS. Local ropivacaine delivery was reported as the amount of ropivacaine extracted per probe. For time-course studies, SPME sampling was performed at the indicated post-treatment intervals to assess the duration of local ropivacaine enrichment after SonoRopi + FUS, with contralateral and sham-treated regions serving as within-animal and treatment controls.

### Widefield calcium imaging of visually evoked cortical activity

Widefield calcium imaging was performed to assess visually evoked cortical activity after FUS-triggered SonoRopi uncaging, using an experimental paradigm adapted from prior widefield GCaMP imaging methods, including ultrasound-compatible dual-channel imaging from the Butts Pauly laboratory (*29*, *30*). Adult Thy1-GCaMP6s mice were used for calcium imaging. To prepare mice for imaging, anesthesia was induced with 4% isoflurane and maintained with 1-2% isoflurane. The scalp and periosteum above the cortex were removed and then a thin layer of Metabond dental adhesive (Parkell) was applied to the skull surface and allowed to dry. Throughout the surgical procedure body temperature was maintained with a heating pad. Mice received 1 mg/kg buprenorphine SR subcutaneously before surgery, and 5 mg/kg carprofen subcutaneously after surgery and daily for three days following surgery. Mice were given at least one week to recover from surgery before imaging experiments. Because these genetically modified mice were used for repeated visually evoked response experiments, a subset of animals contributed to more than one VEP-related condition after a minimum 2-week interval. Reuse was limited to distinct stimulation or treatment conditions; no mouse was re-exposed to the same stimulation condition, and mice that received SonoRopi were not used in subsequent experiments. Each VEP-related experimental condition was repeated in six mice. An isoflurane-control experiment was performed to confirm that 1% maintenance isoflurane caused minimal deviation from baseline visually evoked responses, whereas higher isoflurane concentrations produced greater response deviations (Fig. S1). For treatment experiments, mice were induced with 4% isoflurane for 3 min and then maintained under 1% isoflurane during imaging. Baseline visually evoked responses were recorded before treatment. Mice then received tail vein infusion of SonoRopi or vehicle at a ropivacaine-equivalent dose of 2.5 mg/kg over 5 min. FUS was applied using a 250 kHz transducer targeted to either the right V1 or an off-target left mPFC control site during the final 2.5 min of the infusion. Ultrasound was delivered at an estimated in situ peak negative pressure of 1.1 MPa with a 25% duty cycle using 50 ms pulses repeated every 200 ms.

Visual responses were evoked by stimulation with a flashing white LED light targeted to the eye contralateral to the ultrasound-targeted cortex. Each visual stimulation trial consisted of a 15 s pre-stimulus baseline period, a 5 s flash stimulation period delivered at 10 Hz, and a 10 s post-stimulus period. This stimulation paradigm was repeated 10 times for each imaging session. Widefield GCaMP fluorescence was acquired over the dorsal cortex at baseline and at 5, 30, and 60 min after treatment.

Widefield fluorescence imaging was performed with a PCO.edge 5.5 CMOS camera. Fluorescence excitation was provided by 470 nm and 405 nm LEDs from Thorlabs. The 470 nm calcium-dependent excitation channel was filtered with a 470 nm/40 nm bandpass filter, and the 405 nm reference channel was filtered with a 405 nm/10 nm bandpass filter. The two excitation paths were combined using a 435 nm cutoff dichroic mirror and reflected to the brain using a 495 nm cutoff dichroic mirror, which transmitted green fluorescence emission from the brain to the camera. Emitted fluorescence was filtered with a 525 nm/45 nm bandpass filter before detection. Images were acquired at an overall frame rate of 20 Hz, corresponding to 10 Hz per channel for interleaved 470 nm and 405 nm excitation. The optical configuration followed previously published widefield imaging setups (*29*, *30*). A 3D-printed shielding cone was used to minimize the amount of excitation light reaching the mouse’s eyes and visual stimulus trials were started 30 seconds after the onset of 470/405 nm strobing to allow visual response to the onset of the excitation light to settle back to baseline.

The 470 nm channel was used as the calcium-dependent GCaMP excitation channel, and the 405 nm channel was used as the reference channel for brain hemodynamic and motion-related correction. Image sequences were processed using a dual-wavelength normalization pipeline adapted from prior widefield GCaMP analyses (*29*, *30*). For each trial, ΔF/F was defined as (Fstim − Fpre)/Fpre, where Fstim is the fluorescence signal during visual stimulation and Fpre is the mean fluorescence signal during the pre-stimulus period. Regions of interest were manually defined over right V1 based on anatomical landmarks and visually evoked activation maps. Stimulus-locked ΔF/F traces were averaged across the 10 repeated trials for each animal and time point. Response amplitude was quantified as peak ΔF/F during the 5 s visual stimulation period, and response magnitude was quantified as the area under the ΔF/F curve during the same stimulation period. Treatment-induced changes were compared with pretreatment baseline responses within each animal and across treatment groups.

### S1HL-targeted behavioral sensory testing

Behavioral sensory testing was performed in adult Sprague-Dawley rats to determine whether FUS-triggered SonoRopi uncaging in S1HL produced lateralized changes in hindlimb sensation. Rats were assigned to FUS alone, SonoRopi alone, or SonoRopi + FUS groups. SonoRopi was administered at a ropivacaine-equivalent dose of 2 mg/kg by a 5 min intravenous infusion, and control animals received the corresponding vehicle or sham sonication condition. FUS was targeted to the right S1HL during the final 2.5 min of infusion using the same 250 kHz ultrasound parameters used for in vivo uncaging experiments: 1.1 MPa estimated in situ peak negative pressure and 25% duty cycle, delivered as 50 ms pulses repeated every 200 ms. Because right S1HL represents the contralateral hindlimb, behavioral responses were measured primarily from the left hindlimb, with the right hindlimb serving as an ipsilateral within-animal control.

Mechanical sensitivity was assessed using von Frey filaments and the Dixon up-down method as adapted for rodent paw withdrawal threshold testing (*31*, *32*). Rats were placed on an elevated mesh platform and allowed to acclimate before testing. Calibrated von Frey filaments were applied to the plantar surface of the hind paw, and brisk paw withdrawal, flinching, or licking was scored as a positive response. The next filament force was selected according to the up-down sequence, and the 50% paw withdrawal threshold was calculated from the response pattern. Measurements exceeding the largest tested filament were assigned a ceiling value of 350 g for analysis (*33*, *34*). Mechanical thresholds were measured at baseline and at the indicated post-treatment time points after S1HL-targeted FUS.

Thermal sensitivity was assessed using a unilateral 50 °C hot-plate assay adapted from published procedures (*35*). Rats were gently restrained so that the plantar surface of one hind paw contacted the heated plate, allowing withdrawal latency to be measured separately for each hindlimb. Five measurements were obtained for each hindlimb at 2.5-min intervals and averaged for analysis. A maximum cutoff latency of 20 s was used to prevent tissue injury. Baseline thermal withdrawal latency was measured before treatment, and post-treatment responses were expressed as the change from the corresponding baseline value at the time points shown in Fig. 4D. Withdrawal latencies were analyzed separately for left and right hindlimbs to determine the lateralization, magnitude, and reversibility of the sensory effect.

### Histological safety analysis after brain SonoRopi uncaging

Histological safety after FUS-triggered SonoRopi uncaging was assessed in adult Sprague-Dawley rats using a protocol adapted from our prior AAL safety studies (*18*). Rats were assigned to sham saline, FUS alone, or SonoRopi + FUS groups. SonoRopi was administered at a ropivacaine-equivalent dose of 2 mg/kg by a 5-min intravenous infusion, and FUS or sham treatment was targeted to the right S1HL during the final 2.5 min of infusion using the same sonication parameters used for in vivo uncaging and sensory modulation experiments. At 72 h after treatment, animals were deeply anesthetized and transcardially perfused with 1× PBS followed by 4% paraformaldehyde (PFA) in PBS. Brains were removed, post-fixed in 4% PFA for 24 h, cryoprotected sequentially in 15% sucrose for 48 h and 30% sucrose for 48 h, washed in PBS, and frozen in embedding medium. Coronal sections were cut at 30 μm using a Leica CM1800 cryostat and stored at −20 °C in 0.1 M phosphate buffer containing 30% sucrose and 30% ethylene glycol until staining.

Every 12th section was selected for analysis, corresponding to 360-μm spacing between adjacent analyzed sections. Sections were processed free-floating for hematoxylin and eosin (H&E; Vector Laboratories) to evaluate parenchymal injury and tissue morphology, Fluoro-Jade C (Biosensis) to evaluate neurodegeneration, anti-GFAP immunostaining (Abcam, ab33922) to assess astrocytic activation and gliosis, or anti-IBA-1 immunostaining (Abcam, ab178846) to assess microglial activation. GFAP and IBA-1 primary antibodies were detected with Alexa Fluor 488- and Cy5-conjugated secondary antibodies, respectively (Thermo Fisher Scientific). Stained sections were mounted on microscope slides, and all images were acquired using a Keyence BZ-X800 fluorescence microscope. For quantifiable markers, background-based thresholds and manually defined regions of interest were applied using Keyence BZ-X Advanced Analysis Software.

### Open-field behavioral assessment

Open-field behavior was assessed after right S1HL-targeted treatment using the same infusion and FUS timing used for the histological safety cohort. Rats were assigned to sham, FUS-alone, SonoRopi-alone, or SonoRopi + FUS groups and were placed in a 90 cm × 90 cm open-field arena 15 min after treatment. Behavior was recorded from above for 15 min using a Logitech webcam. Movement tracks were generated from the animal centroid after excluding frames in which tracking failed or the animal was not visible. Total distance traveled was calculated as the cumulative path length during the recording period, and center-time behavior was calculated as the percentage of recording time spent in a predefined 45 cm × 45 cm center zone. Open-field endpoints were analyzed across sham, FUS-alone, SonoRopi-alone, and SonoRopi + FUS groups to assess gross locomotor activity and center-time behavior after treatment.

### Statistical analysis

Statistical analyses were performed using GraphPad Prism version 10. Data are presented as mean ± SD or mean ± SEM as indicated in the figure legends. Except for pharmacokinetic parameter comparisons in Table S1, statistical comparisons were performed using one-way or two-way analysis of variance (ANOVA), as appropriate for the experimental design, followed by Tukey correction for multiple comparisons where applicable. For longitudinal imaging or behavioral datasets, repeated-measures two-way ANOVA was used when the same animals were measured across time. Pharmacokinetic parameters in Table S1 were compared between formulations using Welch unpaired t tests. Exact n values, statistical tests, correction methods, and P values are reported in the figure legends or source data. P < 0.05 was considered statistically significant.

## Acknowledgments

We would like to thank Ronald Watkins for his assistance in producing the ultrasound hardware used in these experiments. We would also like to thank anesthesiologists Tim Meakem MD, Andrew Neice MD, Michael Bokoch MD, PhD, Ben Deverett MD PhD, Vivianne Tawfik MD PhD, and Boris Heifets MD, PhD for their clinical perspective. Additionally, we thank all previous and current members of the Airan and Butts Pauly Labs for their helpful discussions and support.

## Funding

Seed Grant from the Stanford Wu Tsai Neurosciences Institute (RDA)

High Risk Award from the Focused Ultrasound Foundation (RDA)

NIH BRAIN Initiative (NIH/NIMH RF1MH114252, NIH/NINDS UG3NS114438 to RDA)

NIH HEAL Initiative (NIH/NINDS UG3NS115637 to RDA)

Anonymous Donor to the Stanford SOM Radiology Department (RDA)

## Author Contributions

Conceptualization: YX, PJM, RDA

Methodology: YX, PJM, KSR, MLP, MPP

Investigation: YX, PJM, KSR, MLP, MPP, BJY, ARH, MMA, TK, LA, EC, RR, RS

Visualization: YX, PJM

Funding Acquisition: RDA

Project Administration: RDA

Supervision: KBP, RDA

Writing—original draft: YX, PJM

Writing—review & editing: YX, PJM, KSR, MLP, MPP, BJY, ARH, MMA, TK, LA, EC, RR, RS, KBP, RDA

## Data Availability

Source data and code used for analysis will be made available upon publication.

