## Supplemental Materials for "Ultrasonic local anesthetic uncaging enables noninvasive, targeted, and transient knock-out of brain function"

**This file includes:**

Supplementary Methods

Figs. S1 and S2

Tables S1 to S5

Supplementary Methods

**Cryo-electron microscopy (cryo-EM)**

Cryo-EM was performed using a Thermo Fisher Scientific Glacios transmission electron microscope operating at 200 kV. Purified SonoRopi samples (3 μL) were applied to glow-discharged holey carbon grids (Quantifoil Cu R1.2/1.3, 200 mesh). Grids were glow-discharged for 40 s at 15 mA using a PELCO easiGlow system, blotted in a Vitrobot Mark IV for 3 s with blot force 2 at 22°C and 100% humidity, and plunge-frozen in liquid ethane. Frozen grids were clipped, loaded into the microscope, and imaged at 36,000× magnification. Images were recorded using SerialEM software and a K3 direct electron camera with a pixel size of 1.15 Å. Autofocusing was performed with a defocus range of 1.5 to 3 μm. Particle diameters were quantified from cryo-EM images by measuring individual vesicles and plotting count ratios by diameter bin.

**High-performance liquid chromatography (HPLC)**

HPLC analysis was performed on an Agilent 1260 Infinity II HPLC system equipped with a multiple-wavelength UV detector (G7165A) and Agilent OpenLab CDS software. Separation was performed at ambient room temperature using an Agilent 5 HC-C18(2) column (250 mm × 4.6 mm, 5 μm particle size; cat. no. 588905-902) with an Agilent 5 HC-C18(2) guard column (12.5 mm × 4.6 mm, 5 μm; cat. no. 520518-904). Mobile phase A was water with 0.1% trifluoroacetic acid, and mobile phase B was acetonitrile with 0.1% trifluoroacetic acid. Samples were eluted using the gradient in Table S2. Ropivacaine was quantified by UV detection at 263 nm using calibration standards prepared from a ropivacaine reference standard.

**Unencapsulated ropivacaine quantification**

For unencapsulated drug analysis, liposomes diluted 1:4 in PBS were loaded onto a PD-10 size-exclusion column. The initial liposome-containing eluted fraction and the subsequent free-drug eluted fraction were collected separately. Ropivacaine concentration in each fraction was determined by HPLC, and the unencapsulated fraction was calculated as the amount of ropivacaine in the free-drug fraction divided by the sum of ropivacaine in the liposome-containing and free-drug fractions.

**LC-MS/MS analysis**

LC-MS/MS analysis was performed on an Agilent 1290 Infinity LC system coupled to an Agilent 6490 triple quadrupole LC/MS system with iFunnel technology. Data were acquired in positive-ion mode using Agilent Jet Stream electrospray ionization and Agilent MassHunter Workstation Data Acquisition software, and data were processed using MassHunter Quantitative Analysis software. The fragmentor voltage was 380 V and the cell accelerator voltage was 5 V for all transitions. Source parameters and MRM transitions are listed in Tables S3 and S4.

Chromatographic separation was performed at room temperature on an Agilent ZORBAX RRHD Eclipse Plus C18 column (5 cm × 2.1 mm, 1.8 μm particle size; part no. 959757-902) equipped with an Agilent ZORBAX RRHD Eclipse Plus C18 guard column (2.1 mm, 1.8 μm; part no. 821725-901). Mobile phase A was LC/MS-grade water with 0.3% formic acid, and mobile phase B was LC/MS-grade acetonitrile with 0.3% formic acid. The injection volume was 5 μL, the flow rate was 0.300 mL/min, the autosampler temperature was 5°C, and the column compartment was set to 25°C. The chromatographic gradient is listed in Table S5. Calibration standards containing ropivacaine and internal standard were prepared in methanol/water (9:1, v/v) containing 1% formic acid. Analytical responses were calculated as analyte-to-internal-standard peak area ratios and converted to extracted amounts using the calibration curve.

**Pharmacokinetic (PK) analysis**

Noncompartmental analysis was performed from the Fig. 2A blood concentration data. C0 was estimated by log-linear back-extrapolation from the first two observed time points, and terminal parameters were estimated using a log-linear fit to the final three time points (30, 40, and 60 min). Clearance and volume terms assume a 2 mg/kg ropivacaine-equivalent intravenous dose and concentration units of ng/100 μL blood; terminal extrapolated parameters should be interpreted cautiously because the main formulation difference occurred early after dosing.

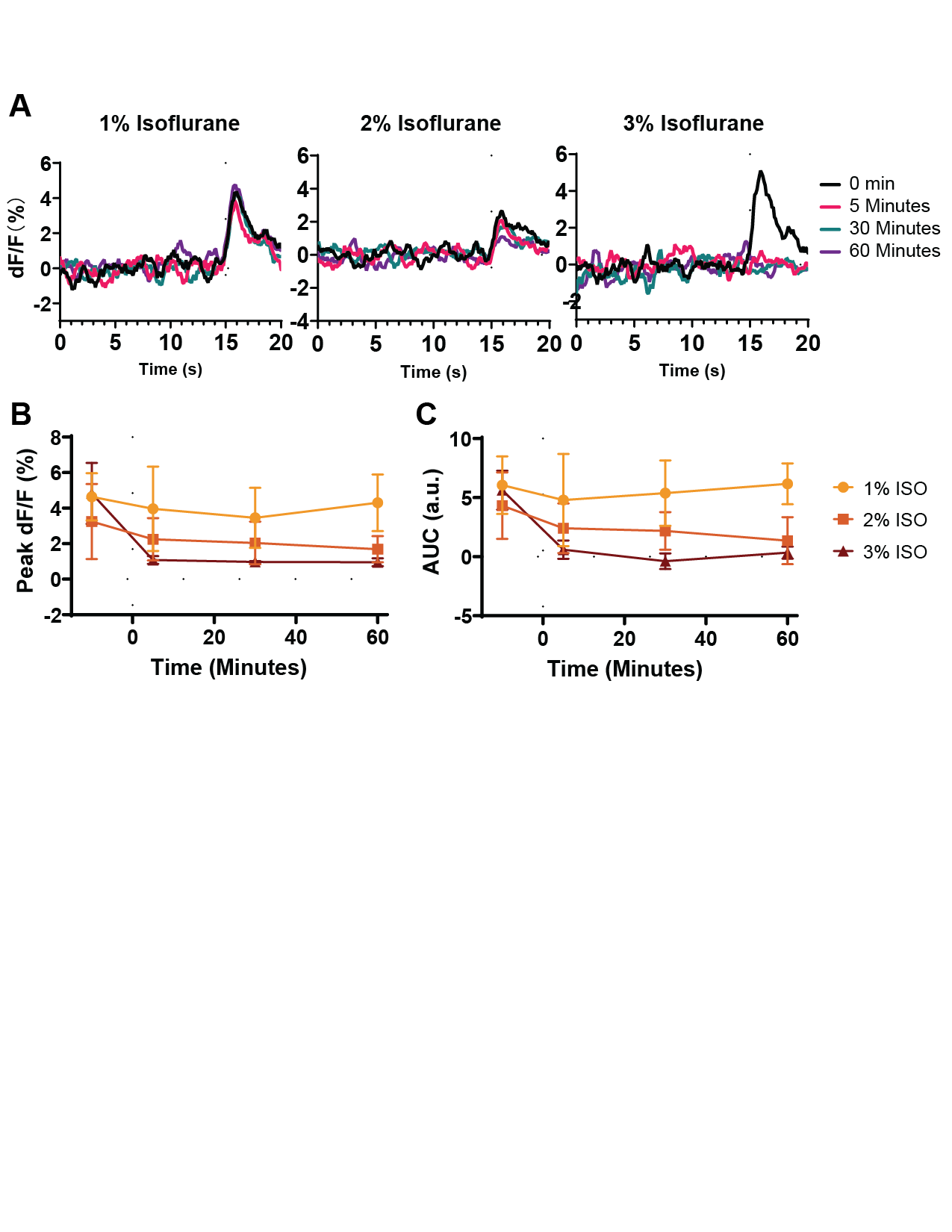

**Fig. S1.** **Effect of maintenance isoflurane concentration on visually evoked cortical responses.** (A) Time course of visually evoked right V1 ΔF/F responses measured at baseline (0 min) and at 5, 30, and 60 min during maintenance with 1%, 2%, or 3% isoflurane. (B) Peak ΔF/F and (C) area under the ΔF/F curve (AUC) of the stimulus-evoked responses at each time point. Peak ΔF/F and AUC remained stable under 1% isoflurane. At 2% isoflurane, both measures showed a progressive downward trend from 5 to 30 and 60 min, whereas 3% isoflurane produced a significant reduction between 5 and 30 min and lasted till 60min. Data are mean ± SEM; n = 3 mice.

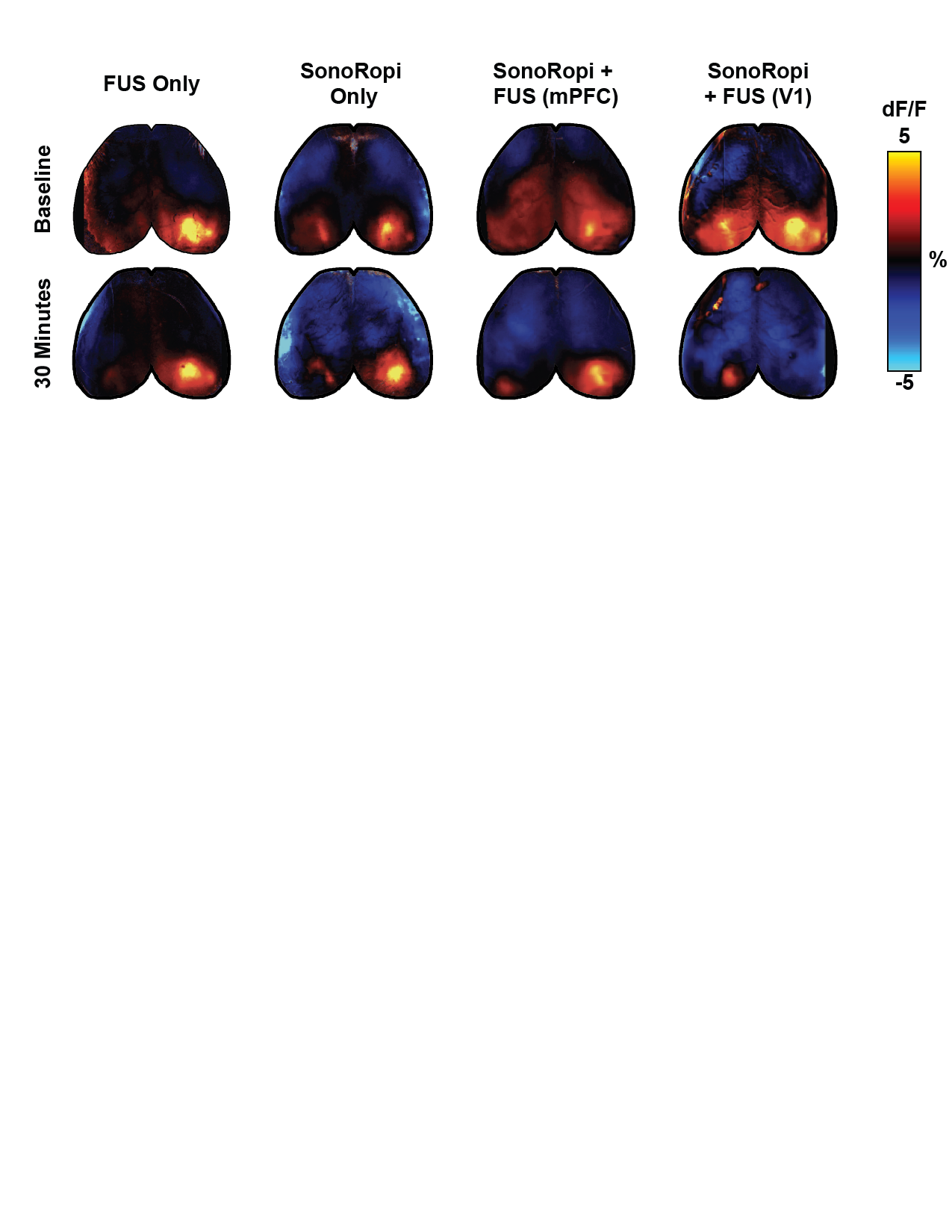

**Fig. S2.** **Representative stimulus-evoked cortical activity maps from individual mice at the pretreatment baseline and 30 min after treatment.** Visually evoked activation in right V1 was reduced at 30 min after V1-targeted SonoRopi + FUS, whereas activation was preserved after FUS alone, SonoRopi alone, and off-target mPFC-targeted SonoRopi + FUS. All maps are displayed using the same ΔF/F color scale.

**Table S1.** Pharmacokinetic analysis of blood ropivacaine after intravenous bolus administration.

| **Parameter** | **Units** | **SonoRopi (mean ± SD)** | **Free ropivacaine (mean ± SD)** | **P value** |
| --- | --- | --- | --- | --- |
| Cmax | ng/100 μL | 151.69 ± 32.48 | 59.03 ± 13.04 | 0.0278 |
| Tmax | min | 1.00 ± 0.00 | 1.00 ± 0.00 | NA |
| C0 | ng/100 μL | 249.55 ± 88.19 | 75.23 ± 21.78 | 0.0698 |
| t1/2 | min | 47.47 ± 10.35 | 147.74 ± 85.00 | 0.0983 |
| AUC0-t | ng/100 μL·min | 1566.13 ± 107.34 | 1305.95 ± 162.65 | 0.0518 |
| AUC0-∞ | ng/100 μL·min | 2241.71 ± 96.37 | 4483.16 ± 1582.31 | 0.0656 |
| MRT0-t | min | 16.85 ± 1.59 | 24.22 ± 1.08 | 0.0043 |
| MRT0-∞ | min | 50.88 ± 12.39 | 204.59 ± 120.06 | 0.0822 |
| CL/D | L/kg/min | 0.089 ± 0.004 | 0.049 ± 0.018 | 0.0176 |
| Vz/D | L/kg | 6.08 ± 1.06 | 8.88 ± 2.40 | 0.1004 |
| Vss/D | L/kg | 4.52 ± 0.95 | 8.50 ± 2.39 | 0.0373 |

Values are mean ± SD. P values are Welch unpaired t tests comparing SonoRopi with free ropivacaine. Cmax, maximum observed concentration; Tmax, time of maximum observed concentration; AUC, area under the concentration-time curve; MRT, mean residence time; CL/D, apparent dose-normalized clearance; Vz/D, apparent dose-normalized terminal volume of distribution; Vss/D, apparent dose-normalized steady-state volume of distribution.

**Table S2.** Chromatographic gradient for HPLC analysis.

| **Time (min)** | **%B** |
| --- | --- |
| 0.00 | 10 |
| 0.30 | 10 |
| 10.30 | 100 |
| 10.50 | 100 |
| 10.75 | 10 |
| 12.00 | 10 |

**Table S3.** LC-MS/MS source parameters.

| **Parameter** | **Setting** |
| --- | --- |
| Source | AJS-ESI, positive mode |
| Gas temperature | 150°C |
| Gas flow | 13 L/min |
| Nebulizer pressure | 30 psi |
| Sheath gas temperature | 375°C |
| Sheath gas flow | 11 L/min |
| Capillary voltage | 2000 V |
| Nozzle voltage | 0 V |
| High-pressure RF | 130 V |
| Low-pressure RF | 75 V |

**Table S4.** Optimized MRM transitions for LC-MS/MS analysis.

| **Compound** | **Internal standard?** | **Precursor ion (m/z)** | **Product ion (m/z)** | **Dwell (ms)** | **Fragmentor (V)** | **Collision energy (V)** | **Cell accelerator (V)** | **Polarity** |
| --- | --- | --- | --- | --- | --- | --- | --- | --- |
| Bupivacaine-d9 | Yes | 298.01 | 149.2 | 60 | 380 | 20 | 5 | Positive |
| Bupivacaine | No | 289.01 | 140.1 | 60 | 380 | 16 | 5 | Positive |
| Bupivacaine | No | 289.01 | 83.9 | 60 | 380 | 40 | 5 | Positive |
| Ropivacaine-d7 | Yes | 282.01 | 136.2 | 60 | 380 | 32 | 5 | Positive |
| Ropivacaine | No | 275.01 | 126.1 | 60 | 380 | 20 | 5 | Positive |
| Ropivacaine | No | 275.01 | 84.1 | 60 | 380 | 56 | 5 | Positive |
| Lidocaine-d10 | Yes | 245.01 | 96.1 | 60 | 380 | 16 | 5 | Positive |
| Lidocaine | No | 235.01 | 86.0 | 60 | 380 | 20 | 5 | Positive |
| Lidocaine | No | 235.01 | 57.9 | 60 | 380 | 36 | 5 | Positive |

**Table S5.** Chromatographic gradient for LC-MS/MS analysis.

| **Time (min)** | **A (%)** | **B (%)** | **Flow (mL/min)** |
| --- | --- | --- | --- |
| 0.00 | 95 | 5 | 0.300 |
| 0.50 | 95 | 5 | 0.300 |
| 4.60 | 5 | 95 | 0.300 |
| 4.90 | 5 | 95 | 0.300 |
| 5.00 | 95 | 5 | 0.300 |
| 5.50 | 95 | 5 | 0.300 |
